# Proteomic Landscape of Pattern Triggered Immunity in the Arabidopsis Leaf Apoplast

**DOI:** 10.1101/2025.02.06.636724

**Authors:** Hsiao-Chun Chen, Carter J. Newton, Gustavo Diaz, Yaochao Zheng, Feng Kong, Yao Yao, Li Yang, Brian H. Kvitko

**Author notes:** Address correspondence to Brian H. Kvitko,. The author responsible for distribution of materials integral to the findings presented in this article in accordance with the policy described in the Instructions for Authors (https://academic.oup.com/plphys/pages/General-Instructions) is Brian H. Kvitko.

## Abstract

The apoplast is a critical interface in plant-pathogen interactions particularly in the context of pattern-triggered immunity (PTI), which is initiated by recognition of microbe-associated molecular patterns (PAMPs). Our study characterizes the proteomic profile of the Arabidopsis apoplast during PTI induced by flg22, a 22 amino acid bacterial flagellin epitope, to elucidate the output of PTI. Apoplastic washing fluid (AWF) was extracted with minimal cytoplasmic contamination for LC-MS/MS analysis. We observed consistent identification of PTI enriched and depleted peptides across replicates with limited correlation between total protein abundance and transcript abundance. We observed topological bias in peptide recovery of receptor-like kinases with peptides predominantly recovered from their ectodomains. Notably, tetraspanin 8, an exosome marker, was enriched in PTI samples. We additionally confirmed increased concentrations of exosomes during PTI. This study enhances our understanding of the proteomic changes in the apoplast during plant immune responses and lays the groundwork for future investigations into the molecular mechanisms of plant defense under recognition of pathogen molecular patterns.

## Introduction

Plants are constantly exposed to microbial pathogens, necessitating the evolution of sophisticated defense mechanisms to ensure survival (Boller and He, 2009; Mott et al., 2014). Pattern-triggered immunity (PTI) represents the first layer of inducible defense, activated by the recognition of conserved microbe-associated molecular patterns (MAMPs) through pattern recognition receptors (PRRs) (Boller and He, 2009). These recognition events trigger a complex regulatory network that initiates a variety of defense responses, including the production of reactive oxygen species (ROS), transcriptional reprogramming, and callose deposition (Mott et al., 2014; DeFalco and Zipfel, 2021). PTI serves as a critical barrier to pathogen invasion and underscores the importance of studying the molecular mechanisms that underpin this process (Boller and He, 2009; DeFalco and Zipfel, 2021).

The apoplast is the intercellular space that contains gas and water, situated between cell membranes and within cell wall matrix (Farvardin et al., 2020). This compartment serves as a critical battleground for plant-microbe interactions and is the location of colonization by foliar bacteria and many other pathogens (Roussin-Léveillée et al., 2024). The apoplast of plants is typically characterized by limited water and nutrient availability, presenting a challenging environment for microbial proliferation (Freeman and Beattie, 2009; O’Leary et al., 2016; Xin et al., 2016; Aung et al., 2018; Gentzel et al., 2022; Liu et al., 2022; Lovelace et al., 2022). The alteration of the apoplastic environment has been proposed to impact pathogen resistance (Roussin-Léveillée et al., 2024). Studies of apoplastic secreted proteins, peptides, and specialized metabolites have provided some insights into apoplastic defense (Anderson et al., 2014; Delaunois et al., 2014; Martínez-González et al., 2018; Gentzel et al., 2022; Serag et al., 2023). Apoplastic proteins have been found to perform diverse functions, including reinforcing the plant cell wall, signal transduction, and inhibition of microbial growth (Alexandersson et al., 2013; Munzert and Engelsdorf, 2025). Secreted proteins include Pathogenesis-related proteins (PRs), enzymes for cell wall modification, enzymes for generation of ROS and redox regulatory proteins (van Loon et al., 2006; Camejo et al., 2016; Nishimura, 2016). Secreted proteases, chitinases, and other hydrolytic enzymes can mediate defense by modification of plant or pathogen structural targets, or the virulence factor targets such as pathogen effector proteins (van der Hoorn, 2008). In addition to secreted proteins, plant extracellular vesicles (P-EVs) are emerging as crucial players in immunity. P-EVs are membrane-bound nanostructures classified into MVBs (multivesicular bodies), EXPO (exocyst-positive organelle), Penetration 1 (Pen1)-positive EVs, vacuoles, and autophagosomes based on their biogenesis (Nemati et al., 2022). They facilitate intercellular and interkingdom communication by transporting bioactive molecules, including proteins and RNAs (Rutter and Innes, 2017; Cai et al., 2019). EVs are implicated in enhancing plant defense through the delivery of immune-related cargo, such as small RNAs and proteins, and contribute to systemic immune signaling (Liu et al., 2021). Exosomes, a type of MVB-derived EV with a median size of approximately 30-150 nm in *Arabidopsis*, have been characterized as a major subtype of P-EVs (Huang et al., 2021). Notably, markers such as tetraspanin-8 (TET8) have been identified in defense-associated exosomes, highlighting their potential role in coordinating immune responses (Wang et al., 2023).

Proteomics approaches using suspension cell culture systems have provided important information of extracellular proteins for understanding the molecular mechanisms involved in pathogen infections (Kaffarnik et al., 2009; Kim et al., 2009). While suspension cell culture systems offer the advantage of limiting cytoplasmic contamination, they do not fully replicate the actual conditions of infection or the specific cellular responses observed in intact tissue apoplast, which are critical for understanding the spatial and temporal dynamics upon pathogens interact. Characterization of apoplastic proteomics typically involve isolating apoplastic washing fluid (AWF), followed by protein identification using techniques such as liquid chromatography-tandem mass spectrometry (LC-MS/MS) (Jung et al., 2008). Isolation of AWF is typically conducted using vacuum or pressure infiltration followed by low-speed centrifugation (Agrawal et al., 2010). This technique is efficient in extracting leaf apoplast contents while minimizing cytoplasmic contamination. To evaluate potential contamination from cytoplasmic components, several assessments are employed using enzymatic assay or immunoblotting to ensure the purity of the AWF (Delaunois et al., 2013; O’Leary et al., 2016). The dynamic nature of the apoplastic proteome and its response to MAMPs remains an active area of research. The Arabidopsis apoplastic proteomic profile during PTI is not well described. Further investigation into the apoplastic proteome under pre-activated PTI will provide crucial insights into the complex interplay of antimicrobial barrier, ultimately leading to the development of more effective strategies for disease control.

In this study, we report the apoplastic proteome of *Arabidopsis thaliana* during PTI induced by flg22 a twenty-two amino acid peptide epitope derived from bacterial flagellin a well-studied MAMP. AWF was extracted by low-speed centrifugation with minimal cytoplasmic contamination and analyzed using LC-MS/MS. Our analyses identified proteins significantly enriched or depleted during PTI. We also compared these proteome profiles with publicly available transcriptomics time-course data, providing insights into the dynamics of early and late PTI outputs. Notably, we observed an increase during PTI in the exosome marker tetraspanin 8 and an increase of exosomes based on nanoflow cytometry. This study provides a detailed snapshot of the proteomic changes in the *A. thaliana* apoplast during PTI elicited by flg22, offering a foundation for understanding the molecular mechanisms governing plant immune responses.

## Results

### Apoplastic washing fluid isolation

A total of 130–150 leaves were collected for apoplastic washing fluid (AWF) isolation from each sample. 3.5–5.0 mL of AWF was recovered per sample across both mock and flg22 treatments. Cytoplasmic contamination was assessed by measuring glucose-6-phosphate dehydrogenase (G6PDH) activity in the AWF, compared to total leaf extracts (Figure S1). The average G6PDH activity in the mock treatment was 0.55 mU/mL, and in the flg22 treatment was 0.16 mU/mL, compared to 14.6 mU/mL in the total leaf extract. Both mock and flg22 samples exhibited <5% G6PDH activity relative to the total leaf extract with no statistic difference between mock and flg22 treatments, indicating minimal cytoplasmic contamination.

### Principal Component Analysis (PCA) of flg22 and mock samples

The overall distribution of log-transformed total protein abundance per sample showed consistency across replicates. Median log protein abundances were similar between the flg22-treated and mock-treated samples, indicating a reliable dataset for further multivariate analysis (Figure S2). Principal Component Analysis (PCA) was used to examine the proteomic differences between flg22-treated and mock-treated samples (Figure 1). The first two components, Dim1 and Dim2, explained 27.9% and 25.9% of the total variance, respectively. The PCA plot revealed clustering based on sample types.

**Figure 1.**
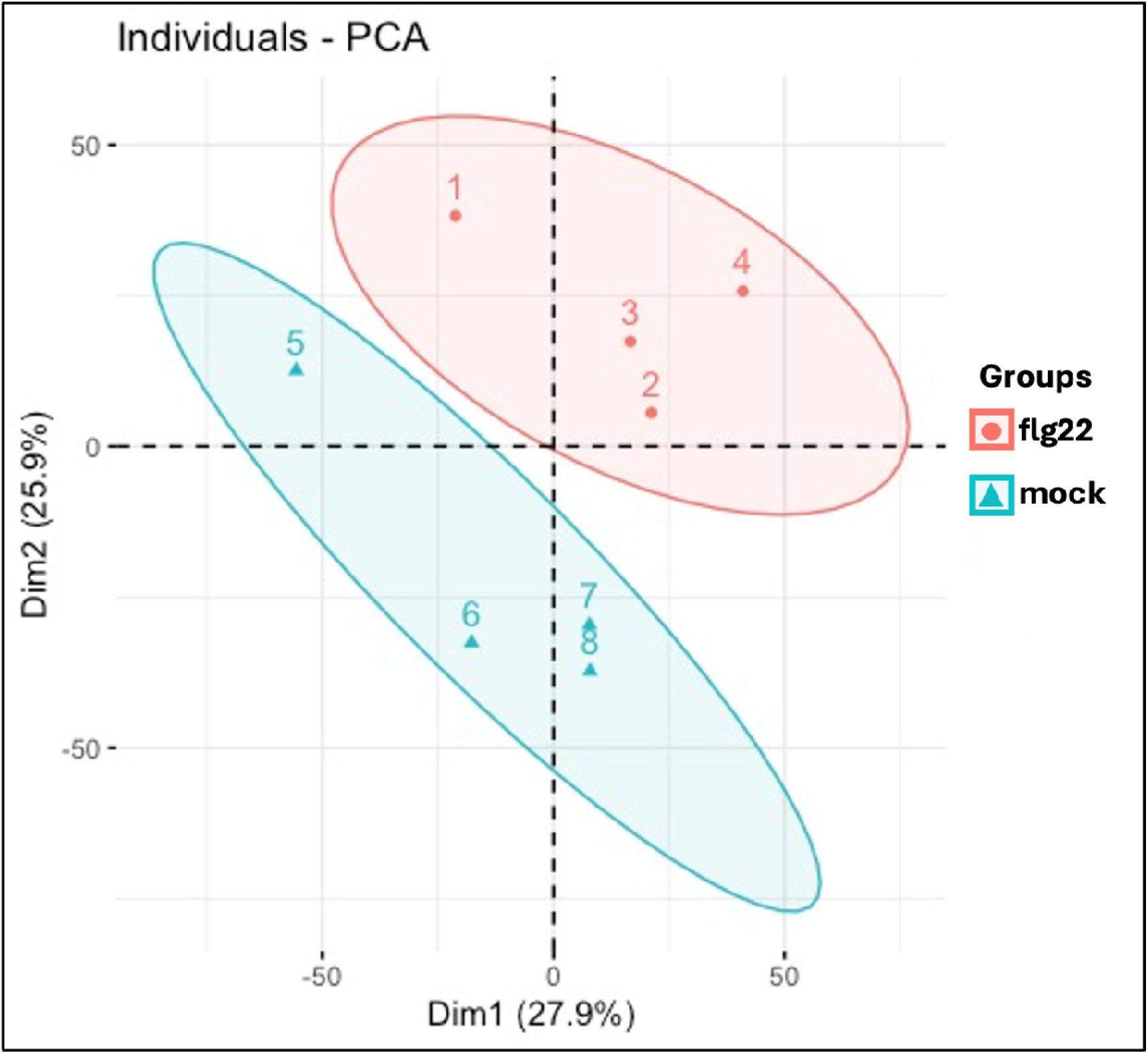
Principal Component Analysis (PCA) of flg22 and mock samples. The PCA plot illustrates the distinct clustering of samples based on their treatment conditions. Flg22-treated samples and mock-treated samples form separate clusters, indicating clear differences in their protein abundance profiles. The first principal component (PC1) accounts for 27.9% of the total variance, while the second principal component (PC2) explains 25.9% of the variance. Each data point represents an individual biological replicate, with the spatial distribution of points within each cluster indicating the degree of variability among replicates within the same treatment group.

### Comparison of PTI apoplastic proteins enrichment relative to time course transcriptome profiles

A total of 108 proteins were identified as significantly enriched in samples at 16 hours post flg22 treatment comparing to mock controls (Log_2_ Fold Change (Log2FC) ≥ 1 and p-value ≤ 0.05) (Figure 2). We manually classified these proteins into six groups based on protein annotations in the open-access information portal Araport: (1) RLKs/RLPs (Receptor-Like Kinases/Receptor-Like Proteins), (2) redox and redox-associated proteins, (3) hydrolytic enzymes, (4) antimicrobial peptides, (5) extracellular vesicle-associated proteins, and (6) others. To better understand to what degree transcriptional regulation relates to protein abundance we integrated our proteomic data with a published Arabidopsis transcriptomics time course from by Hillmer et al., 2017. Within each category, proteins were ranked by their log2FC from highest to lowest abundance. IOS1 and SIF2 were among the most abundant in the RLK/RLP category. Other immune-associated RLKs like RLK902 and several CRKs (Cysteine-rich receptor-like protein kinases) were also enriched (Gao et al., 2024). A considerable number of redox-related proteins, such as peroxidases (PERs) and glutathione-S-transferases (GSTFs) were prominent. Several hydrolytic enzymes were identified in the apoplast during flg22-induced PTI in *Arabidopsis*. Enzymes involved in carbohydrate metabolism, including chitinase family proteins (At1g02360), and beta-hexosaminidase 3 (HEXO3), were enriched. Subtilisin-like protease SBT3.3 and glucan endo-1,3-beta-glucosidase 14 (At2g27500) were also identified, along with related beta-glucosidases. Enzymes involved in cell wall modifications, such as pectin methyl-esterases (PMEs) and xyloglucan endotransglucosylases, were present in the enriched profile. EARLI1 gene in Arabidopsis encodes a lipid transfer protein-like protein that plays a crucial role in mediating systemic defense mechanisms, specifically systemic acquired resistance (SAR) and induced systemic resistance (ISR) were also enriched by flg22 (Vlot et al., 2021). Notably, in flg22-enriched profile, three proteins associated with vesicle trafficking were identified as significantly enriched under PTI conditions. The enriched proteins include Syntaxin of Plants-122 (SYP122), a Qa-SNARE which functions in vesicle fusion with the plasma membrane peptidyl-prolyl cis-trans isomerase FKBP15-1 (FKBP15-1), which may contribute to protein folding within vesicles, and tetraspanin-8 (TET8), associated with exosome extracellular vesicle structural integrity (Boavida et al., 2013; Wang et al., 2020; Rubiato et al., 2022).

**Figure 2.**
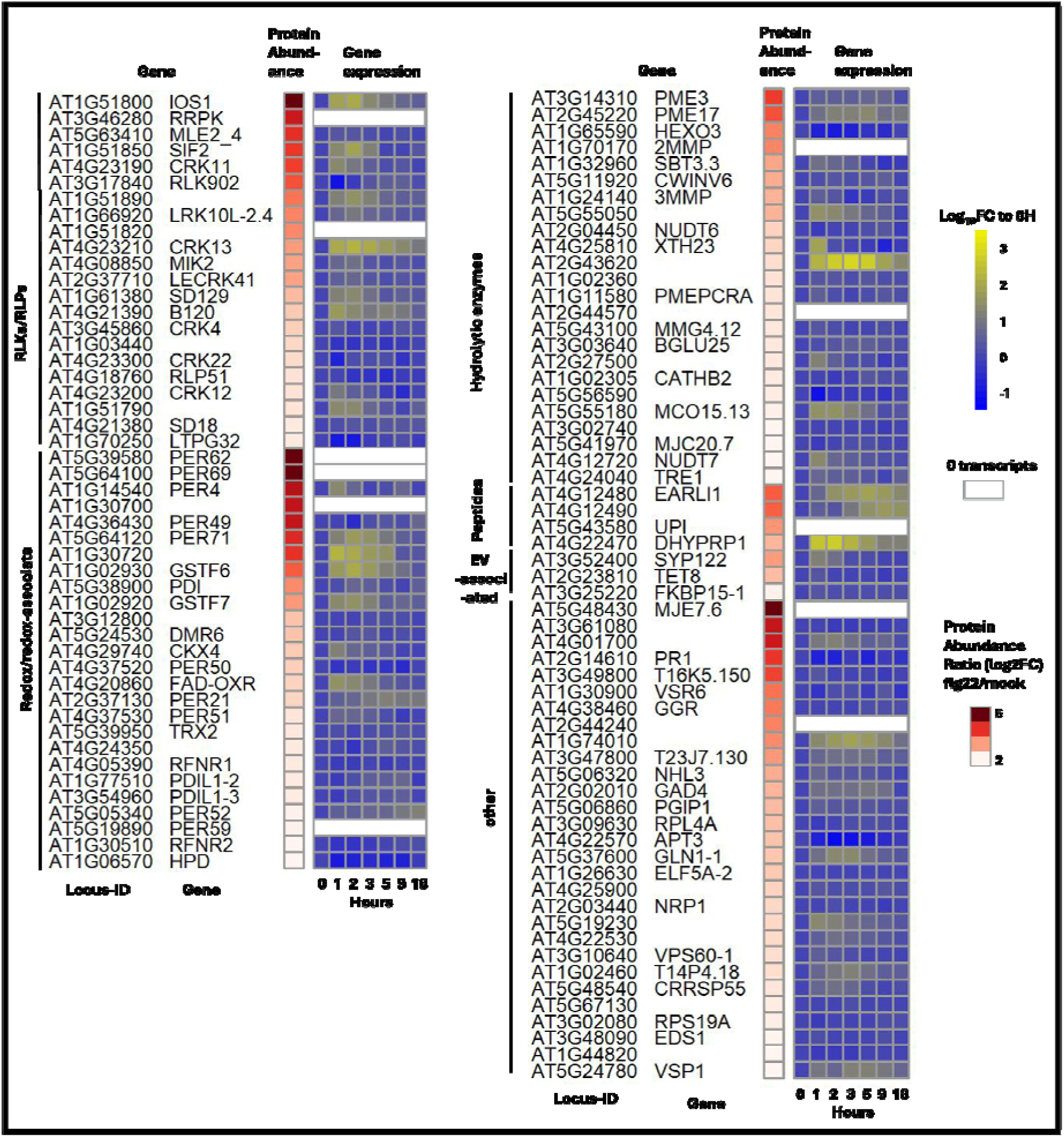
PTI-enriched proteins ranked by functional groups and relative to transcriptome profiles. The heatmap presents flg22-enriched protein abundances on a red scale. The corresponding log10 transformed transcript abundance is displayed on a blue-yellow scale, with data are shown at 0, 1, 2, 3, 5, 9, and 18 hours post flg22 treatment relative to the initial time point (0 hour). Transcriptome data of raw reads were obtained from study of Hillmer et al., 2017. GEO data series GSE78735.

The heatmap (Figure 2) illustrates how the transcript abundance changes over time, providing insights into the continuity or discontinuity between transcription and protein accumulation in response to flg22 treatment. The transcriptomic analysis revealed distinct patterns of gene expression in response to flg22 treatment, corresponding with the functional roles of the identified proteins in PTI. There are 22 receptor-like kinases and receptor-like proteins (RLK/RLP) identified in the enriched profile and 10 of them displayed rapid upregulation at the first hours then the expression level gradually decreased. Several redox-related proteins and hydrolytic enzymes, such as peroxidases (PER21, PER52, and PER71) and Glutathione S-Transferase (GSTF6 and GSTF7) showed sustained transcriptional upregulation at a relatively later time point. Overall, within each functional group, protein abundance ratios did not necessarily align with transcriptional trends in a time-course manner.

### Recovered peptides of RLKs are heavily biased to ectodomains

RLKs membrane proteins were enriched in the apoplastic proteome after flg22 treatment. Considering that these are transmembrane proteins, we would not expect them be found in the free extracellular fraction. We investigated the topological distribution of detected peptides in 18 identified RLKs in our dataset enriched under PTI conditions (Figure 3). This detection pattern shows a notable bias toward the extracellular region, suggesting that portions of these membrane-associated proteins are accessible or released into the apoplast under PTI conditions. Very few peptides were captured close to the cytoplasmic regions. The extracellular bias aligns with the apoplast-specific protein collection procedure. The limited peptide coverage in cytoplasmic regions further emphasize the specificity of the apoplast isolation protocol.

**Figure 3.**
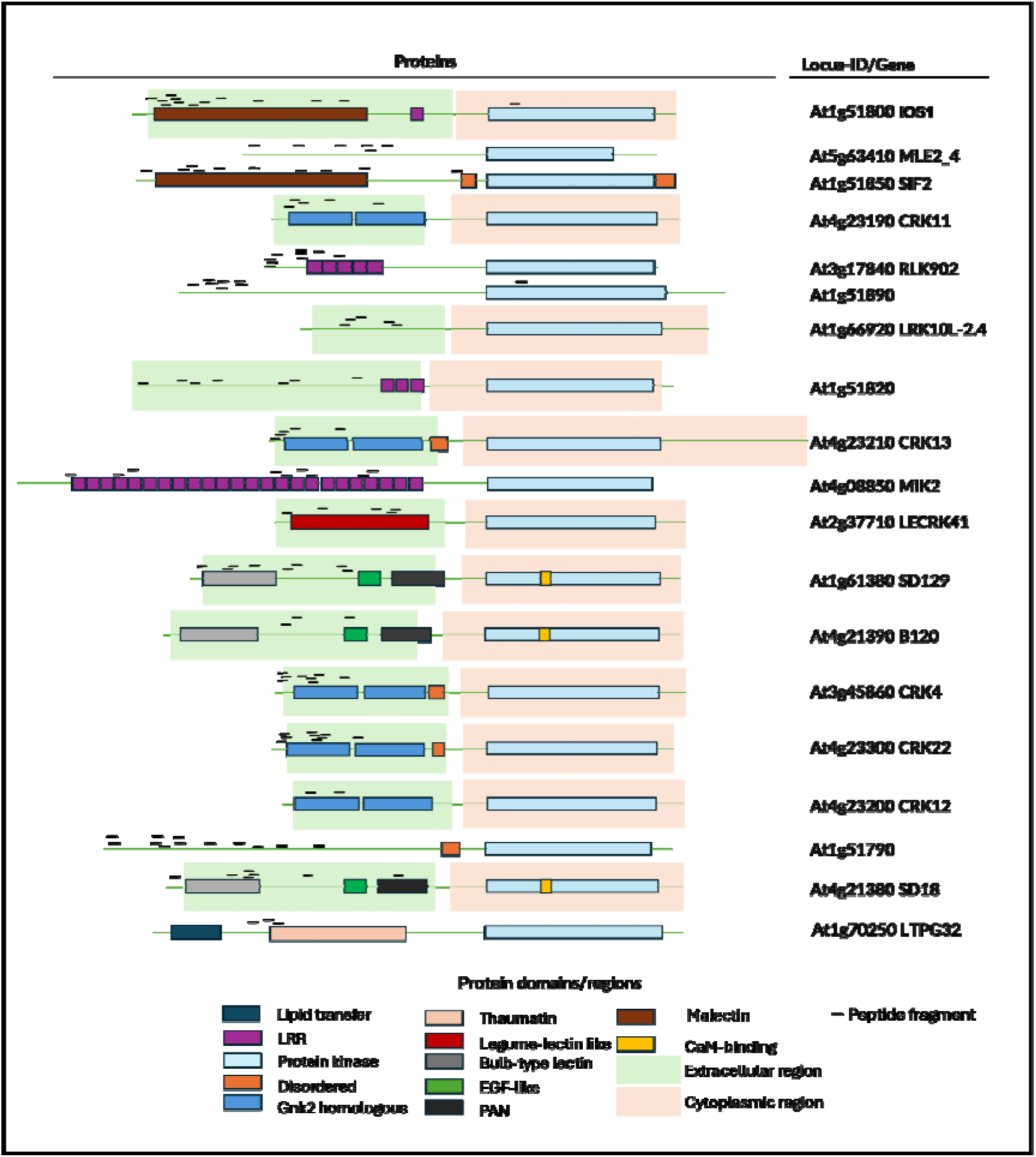
Peptide distribution analysis of receptor-like kinases (RLKs) in flg22-enriched proteins. The schematic illustrates the mapping of identified peptides to PTI-enriched receptor-like kinases (RLKs), represented as short black bars above the protein domain structures. Each RLK is depicted with its extracellular region (green background) and the cytoplasmic kinase region (pink background) on the right. The visualization reveals the majority of detected peptides are localized to the extracellular regions of the RLKs, while the cytoplasmic regions, particularly the kinase domains, show notably fewer peptide matches.

### Apoplastic proteins showing lower abundance by flg22 treatment displayed limited functional clustering and transcript-protein abundance discrepancies

A total of 255 proteins were identified as significantly decreased following flg22 treatment compared to mock conditions ((Log_2_ Fold Change (Log2FC) ≤ −1 and p-value ≤ 0.05) (Figure 4). Unlike the distinct functional clustering observed in the enriched protein profile by flg22 treatment, lower abundant proteins did not exhibit clear patterns linked to PTI or disease response functions. However, based on molecular function GO analysis, these proteins were categorized into 10 functional groups. We observed some genes displaying high transcript levels post-flg22 treatment despite their decreased protein abundance in the apoplast, indicating some discontinuity between transcript and protein abundances.

**Figure 4.**
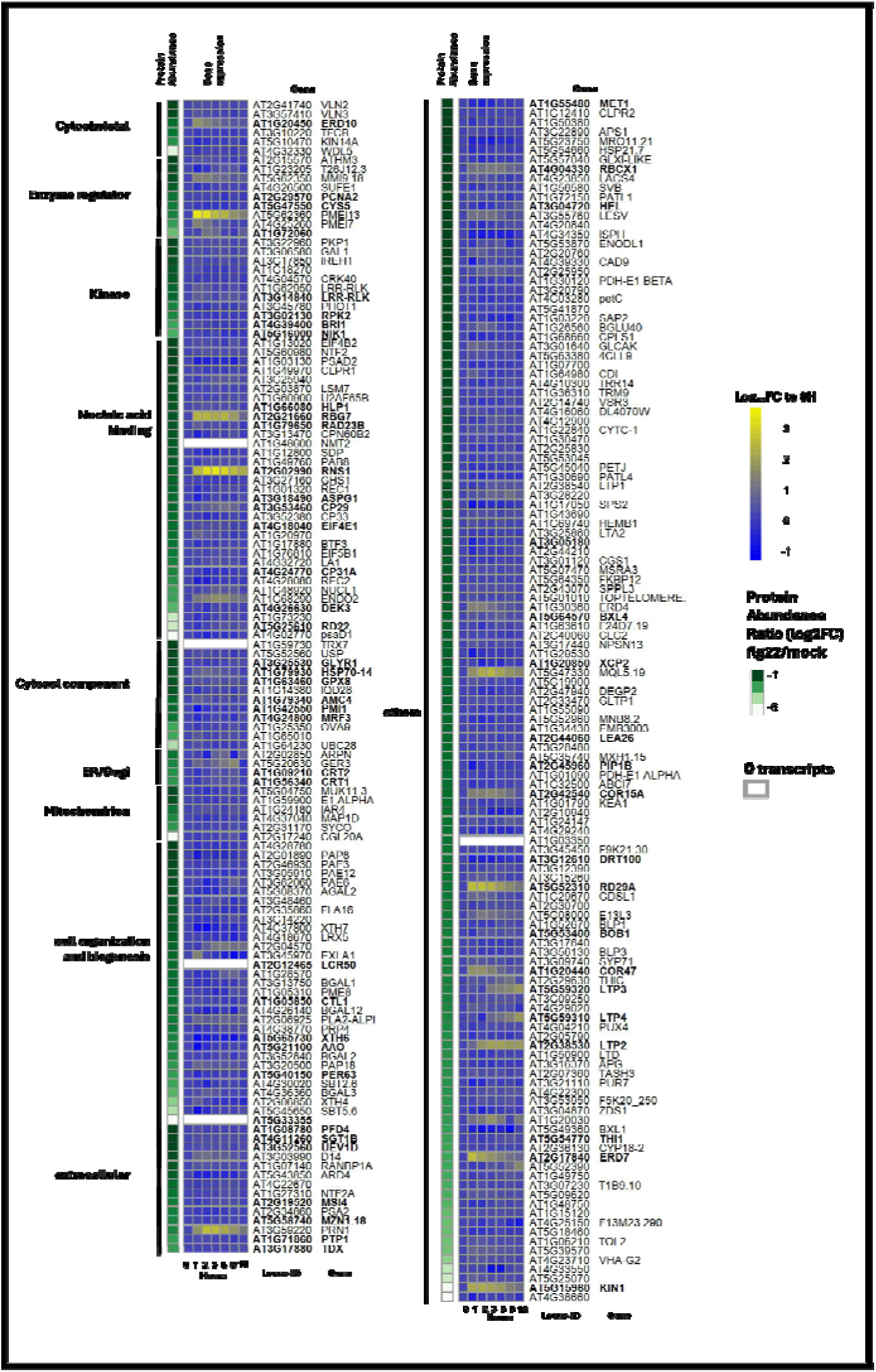
PTI-lower abundant proteins ranked by functional groups and relative to transcriptome profiles. The heatmap illustrates proteins showing lower abundance in the flg22-treated group compared to mock treatment, represented on a green scale. The corresponding log10 transformed transcript abundance is displayed on a blue-yellow scale, with data are shown at 0, 1, 2, 3, 5, 9, and 18 hours post flg22 treatment relative to the initial time point (0 hour). Proteins associated with GO stress responses are highlighted in bold font.

### Gene Ontology (GO) analysis of the enriched protein profile under the flg22 treatment

We performed Gene Ontology (GO) analyses of the 108 significantly enriched proteins in flg22 treatment with TAIR GO Term Enrichment platform. The GO analysis identified significant enrichment of multiple biological processes in flg22-enriched proteins (Figure 5A). Processes with the highest gene number include response to stimulus and response to stress, suggesting that many enriched proteins are involved in both general and specific stress responses. More specific processes, such as response to biotic stimulus, defense response to Gram-negative bacterium, and defense response to fungus, highlight these proteins’ roles in responding to presence of pathogens. Specific processes such as response to oxygen levels and response to oxidative stress are also enriched, suggesting that these proteins may play roles in reactive oxygen species (ROS) production, which are commonly associated with plant immune responses signaling (Torres et al., 2006). Notably, highly representative pathway with higher fold enrichment like defense response to Gram-negative bacterium align with the expected response to the bacterial elicitor flg22.

**Figure 5.**
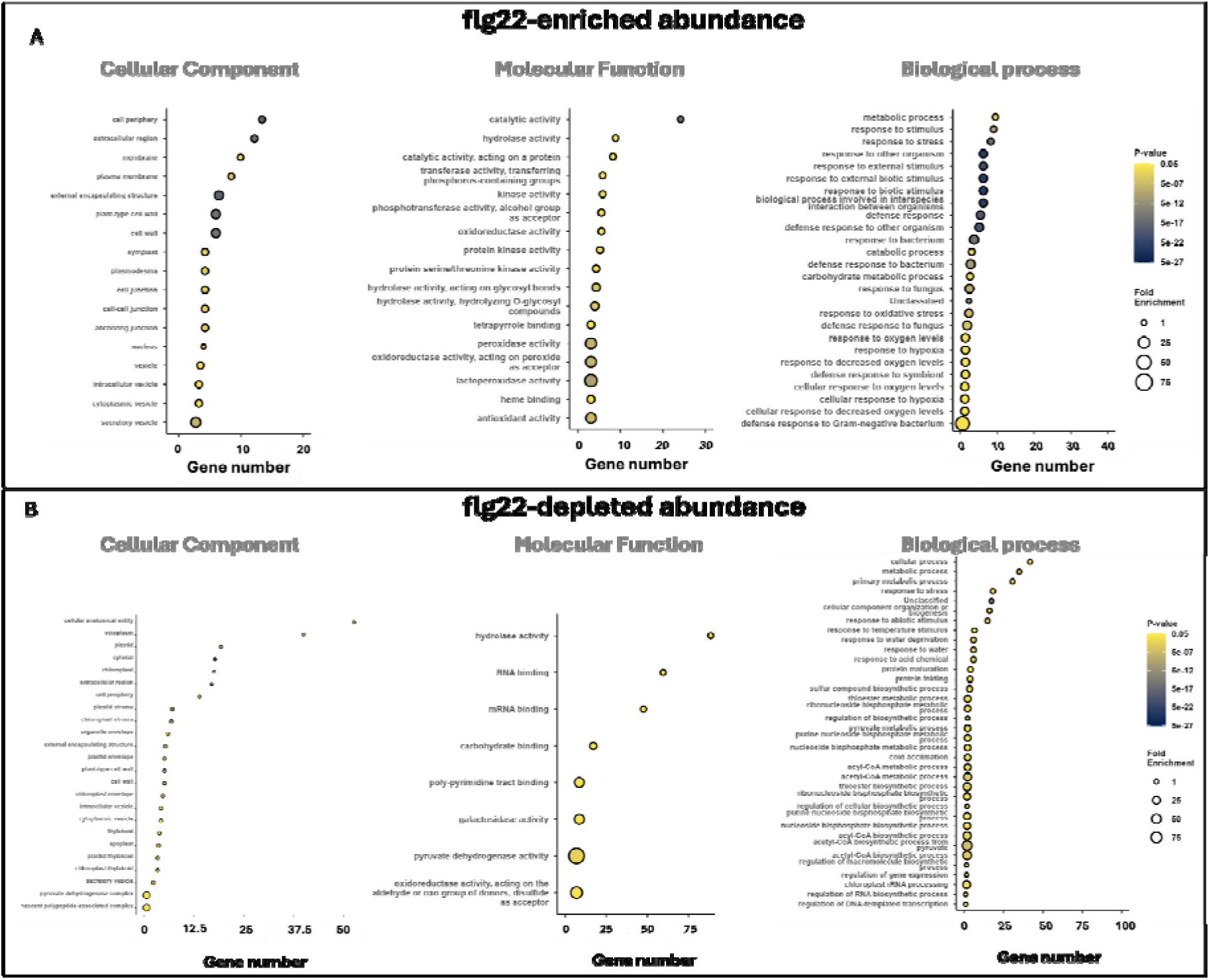
Gene Ontology (GO) analysis of flg22-enriched and flg22-decreased abundance protein profile. Dot plot represents significant GO terms of (A) 108 significantly enriched proteins and (B) 255 significantly decreased proteins categorized into cellular component, molecular function, and biological process. The x-axis represents the gene number of each category, while the size of the circles indicates fold enrichment. Color intensity of the circles corresponds to the statistical significance of enrichment (−log10 adjusted p-value). GO terms are arranged on the y-axis, grouped by their respective ontologies (cellular component, molecular function, and biological process).

The molecular function analysis for flg22-enriched proteins revealed significant enrichment in various catalytic and kinase activities. Nearly 25% of the proteins identified in the GO analysis were annotated for catalytic activity. The enrichment of oxidoreductase activity, including peroxidase and lactoperoxidase functions, suggests roles associated with redox reactions and potential oxidative stress responses. Proteins with kinase and transferase activities, particularly those transferring phosphorus-containing groups, were also prominent. Specifically, protein kinase activity and protein serine/threonine kinase activity were enriched, indicating involvement in phosphorylation cascades.

Consistent with an apoplastic proteome, the cellular component analysis for flg22-enriched proteins revealed significant enrichment in regions associated with the cell periphery, extracellular region, and membrane. Nearly 15% of the proteins identified in the GO analysis were localized to the cell periphery, suggesting a strong role in cell surface interactions. Notably, vesicles and components associated with the secretory pathway were also prominent in the dataset. The presence of proteins localized to vesicles, including secretory and cytoplasmic vesicles, underscores their potential involvement in protein trafficking and secretion.

In contrast to the flg22-enriched proteins, which exhibited distinct functional categories related to pattern-triggered immunity (PTI) and defense responses, the proteins with lower abundance did not display clear patterns linked to these functions (Figure 5B). However, the GO analysis revealed that these lower abundance proteins were involved in a wide range of cellular processes, including metabolism and cellular turnover. The cellular component analysis of these proteins showed their overall localization in the apoplast, with involvement in protein secretion and cell periphery, albeit with small fold enrichment. The involvement of these lower abundance proteins in the secretory pathway underscores their role in collecting extracellular proteins in the apoplast, even if their abundance is reduced.

### PTI is associated with increased numbers of extracellular vesicles (EVs) in leaf apoplast

Several studies have reported the enrichment of plant extracellular vesicles (EVs) during pathogen infection (Rutter and Innes, 2017; Cai et al., 2019). In our dataset, we observed significantly enriched proteins associated with vesicular export, including SYP122, TET8, and FKBP15-1. SYP122 and TET8 show increased change of 8.0 and 5.2-fold, respectively, relative to the mock condition. Notably, TET8 is an exosome marker involved in exosome stability (Cai et al., 2019; Liu et al., 2024). Exosomes are one of several classes of EVs with a size typically between 50nm-150nm in dimeter and have been associated with the cross-kingdom delivery of small RNA for RNAi. We isolated plant EVs using ultracentrifugation and quantified them using Nano-flow cytometry. The analysis of EV in both flg22-treated and mock samples reveals that the median size of EVs in flg22-treated samples is 67.8 nm, compared to 70.7 nm in mock samples (Figure 6A). This similarity in size indicates no significant difference between the two conditions, suggesting that the majority of these vesicles fall within the size range characteristic of plant exosomes (Rutter and Innes, 2017; Cai et al., 2019). Exosomes were found at 5.1 × 10 particles/mL in the flg22-treated samples, nearly three times higher than the 1.7 × 10 particles/mL observed in the mock samples (Figure 6C). This increased concentration in the flg22 condition aligns well with the 5-fold higher protein levels identified in the flg22/mock proteomics analysis. The staining of the AWF-derived EVs with the lipid dye, CFSE, highlighted the shape and size characteristic of exosomes (Figure 6D).

**Figure 6.**
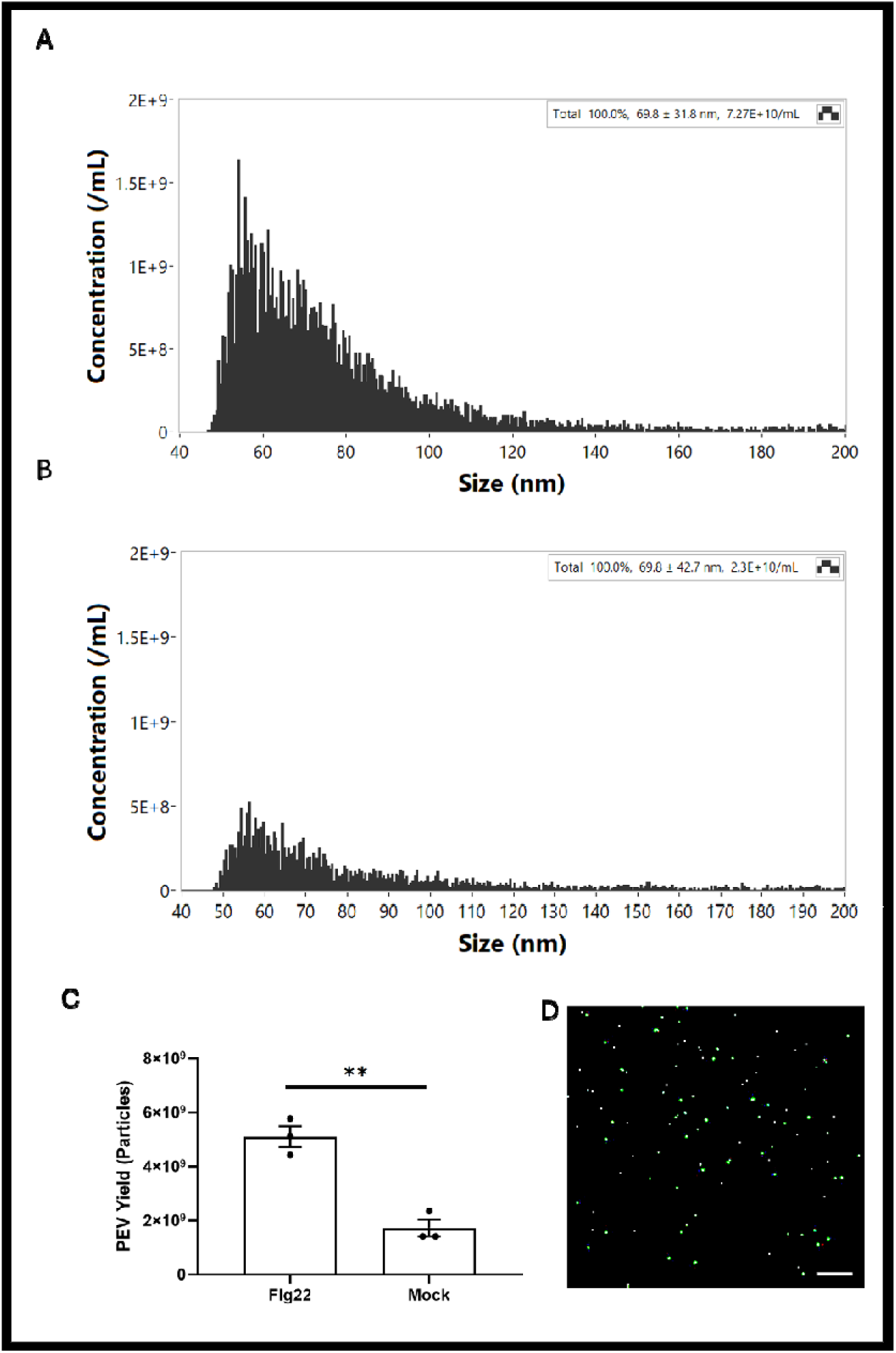
Plant extracellular vesicles are enriched in flg22 treatment. (A) Size distribution of flg22-P-EVs identified by Nano-flow cytometry (NFC). (B) Size distribution of mock-P-EVs identified by Nano-flow cytometry (NFC). (C) Comparison of P-EV production between the flg22 and mock groups, showing a significant increase in P-EV production in the flg22 group. Statistical analysis was performed using two-tailed unpaired Student’s t-test to determine P-EV yield differences between groups with \*\**P* <0.01. (D) Confocal images of CFSE-stained P-EV from mock samples. Scale bar=50μm.

## Discussion

GO analysis of the PTI enriched apoplastic proteome revealed significant enrichment in predicted biological processes such as defense responses to Gram-negative bacteria and oxidative stress. The molecular functions enriched in flg22-treated samples included both kinase and oxidoreductase activities, consistent critical roles of PRRs and redox regulation in plant defense mechanisms. The cellular component GO terms, as would be expected, emphasized the extracellular localization of both enriched and depleted proteins, including the cell wall, plasma membrane, secretory vesicles, and extracellular regions. Unlike the PTI-enriched proteins, which were prominently associated with defense signaling and oxidative stress management, the PTI-depleted proteins displayed functions more broadly related to cellular homeostasis and general secretion mechanisms. This may be due to less cytoplasmic leakage in flg22-treated samples during the collection process (Figure S1) or may reflect an overall shift in molecular function. Notably, more than 18% of depleted proteins were associated with functions related to cell organization and biogenesis, including BRI1 (BRASSINOSTEROID INSENSITIVE 1), indicating the stabilization of BR signaling under naïve conditions, this can potentially be a consequence of the growth-defense trade-off (Huot et al., 2014).

We defined five functional categories of apoplastic proteins enriched during flg22-triggered PTI, including RLK/RLPs, redox-associated proteins, hydrolytic enzymes, antimicrobial peptides, extracellular vesicle (EV) associated proteins. Among the RLKs, IOS1 (Impaired Oomycete Susceptibility 1) and SIF2 (Stress Induced Factor 2) were highly enriched. These two malectin-domain RLKs have both been observed previously to play roles in PTI signaling. IOS1, a malectin-like leucine-rich repeat RLK, modulates Arabidopsis immunity by forming complexes with FLS2 and BAK1 to amplify pathogen recognition and defense responses. Loss-of-function mutants of IOS1 demonstrate increased susceptibility to bacterial pathogens (Yeh et al., 2016). Similarly, SIF2 also interacts with FLS2 and BAK1 and has been implicated in early flg22-triggered immune signaling, contributing to downstream defense activation (Chan et al., 2020). The peptide recovery analysis of RLKs demonstrated a marked bias toward the receptor extracellular regions, with minimal peptides detected in their cytoplasmic regions. The limited recovery of cytoplasmic peptides further aligns with the low levels of cytoplasmic contamination in the apoplastic washing fluid, emphasizing the effectiveness of the apoplast isolation protocol as described by Lovelace et al., 2022. The recovery of free ectodomains in the apoplast may be an artifact of the sampling procedure or could indicate that these ectodomains are endogenously released into the apoplast under the tested conditions through an unknown mechanism.

A comparative analysis of transcriptome and proteome profiles revealed a complex relationship between transcriptional regulation and endpoint protein abundance during flg22-triggered PTI. Several high-abundant proteins displayed consistent trends between transcript levels and protein accumulation over the time course. For example, IOS1 and SIF2, both receptor-like kinases (RLKs), exhibited rapid transcriptional upregulation within the first few hours of flg22 treatment. This aligns with their role in early pathogen recognition and activation of downstream signaling pathways (Bigeard et al., 2015). Similarly, Peroxidases such as PER15 (Peroxidase 15), showed a strong correlation between transcript and protein levels, consistent with its involvement in reactive oxygen species (ROS) management and oxidative stress responses during PTI. However, other high-abundance proteins demonstrated protein enrichment despite declining transcript levels over time. Cysteine-Rich RLKs, another set of RLKs, have distinct patterns of expressions despites their role in response to flg22 perceptions (Yadeta et al., 2017). For instance, CRK11 and CRK12 remained abundant in the apoplast even as their transcript levels decreased after the initial hours of flg22 exposure. On the other hand, CRK13 exhibited decreased transcript levels over time but maintained a high expression pattern and protein abundance during late PTI, suggesting its crucial role in enhanced resistance to the bacterial pathogen *Pseudomonas syringae* (Acharya et al., 2007). These suggested post-transcriptional regulatory mechanisms, such as enhanced mRNA stability or protein stabilization, which might prolong the functional presence of critical immune components. Such discrepancies reflect the likely importance of post-transcriptional regulation and protein turnover dynamics in fine-tuning the immune response.

The enrichment of various hydrolytic enzymes in apoplast in response to flg22 treatment suggested a multifaceted response to PTI. Several enzymes involved in plant cell wall modification were found to be enriched including pectin methyl-esterases (PME3, PME17), xyloglucan endotransglucosylase/hydrolase (XTH23), and β-1,3-glucanases (At2g27500 and At5g56590) (Bethke et al., 2014; Perrot et al., 2022; Zhang et al., 2022). These enzymes collectively suggest that the plant is actively remodeling its cell wall during PTI, possibly to enhance its barrier function against potential pathogens. Several hydrolytic enzymes capable of N-glycosylation of immune receptor and degrading pathogen-associated targets (especially fungal cell wall) were enriched, such as β-hexosaminidase 3 (HEXO3) and endochitinases (At2g43620 and At1g02360) (Liebminger et al., 2011; Fiorin et al., 2018). These enzymes suggest that the plant is actively degrading potential pathogen-derived molecules, which could both weaken a broad-spectrum of pathogens and generate additional immune-stimulating signals. Various proteases and protein-modifying enzymes were also enriched. Metalloendoproteinases (2-MMP and 3-MMP) and subtilisin-like protease (SBT3.3) may be involved in processing defense-related proteins within the extracellular matrix or degrading pathogen-derived proteins (Flinn, 2008; Ramírez et al., 2013; Zhao et al., 2017). Plant aspartyl proteases (MMG4.12 and At3g02740) and cathepsin B-like protease 2 (CATHB2) were also found. These proteases contribute to plant immunity through various mechanisms, from immune signaling activation and systemic resistance to direct cleavage of pathogen proteins (McLellan et al., 2009; Figueiredo et al., 2021). The presence of these proteases suggests active protein processing and turnover, which could be important for the homeostasis of defense signaling molecules and limiting pathogen fitness. Other metabolic enzymes, such as NUDT6 and NUDT7, members of the nudix hydrolase family, were enriched by flg22. They play crucial roles in regulating plant immunity and stress responses by maintaining the redox balance of NADH and acting as ADP-ribose pyrophosphatases (Fonseca and Dong, 2014). The coordinated action of these hydrolytic enzymes with various types of substrates likely contributes to the rapid and effective immune response against potential pathogens, highlighting the sophisticated defense mechanisms employed in pathogen recognition and immune activation.

The observation of pathogenesis-related 1 (PR1) protein enrichment in the flg22 treatment profile, with a notable 4.67-fold change, strongly aligns with key findings in induced pattern-triggered immunity (PTI). PR1 is a well-established marker for salicylic acid (SA)-regulated plant immunity, and its secretion is critical for activating systemic acquired resistance (SAR). Upon flg22 treatment, plants exhibit increased PR1 expression as part of their PTI response (Djamei et al., 2007). The significant enrichment of PR1 protein abundance observed in this study confirms its responsiveness to PTI triggered by flg22 treatment and further demonstrates the effectiveness of the experimental conditions in inducing a robust immune response.

For lower abundant proteins by flg22 treatment, inconsistencies with transcript levels were even more pronounced. Many of these proteins, such as certain housekeeping enzymes and stress-related factors, exhibited little to no transcriptional change over the time course. Their reduced abundance in the apoplast may result from active degradation, inhibited secretion, or preferential retention in intracellular compartments. Unlike high-abundant proteins, these low-abundant factors were associated with less PTI-relevant processes, further indicating that their depletion may reflect cellular reorganization rather than active participation in immune responses. The integration of transcriptomics and proteomics provides additional insights into the temporal coordination of immune responses. Further time-resolved multi-omics studies could elucidate the functional significance of these regulatory discrepancies, revealing additional layers of complexity in plant immunity.

Several extracellular vesicles (EVs)-associated proteins exhibited a notable increase in flg22-treated apoplast samples. In particular, tetraspanin-8 (TET8), a well-established exosome marker, was enriched fourfold compared to mock-treated samples. Cai et al. (2018) previously identified TET8 as a core component of exosomes involved in stress responses, and pathogen defense as mediators of interkingdom RNAi. Additionally, the concentration of EVs in flg22-treated samples was approximately three times higher than in mock-treated samples. This increase in EVs production suggests enhanced trafficking of defense molecules and signaling factors to the apoplast. The enrichment of Syntaxin-122 (SYP122), a SNARE protein involved in vesicle fusion, supports the idea that vesicle-mediated transport is a key component of the immune response (Waghmare et al., 2018). These results underscore the importance of EVs as critical components of PTI, facilitating intercellular communication and the targeted delivery of antimicrobial proteins and signaling molecules. The substantial enrichment of TET8 and the increased exosome particles suggest that EVs play a pivotal role in coordinating and amplifying defense responses during pathogen attack.

Overall, our study provides a snapshot of the apoplastic proteome in *Arabidopsis* during the later phases of flg22-induced pattern-triggered immunity (PTI). A total of 108 significantly enriched proteins were identified and categorized into key functional groups, including receptor-like kinases/receptor-like proteins (RLK/RLPs), redox-related proteins, hydrolytic enzymes, small peptides, and extracellular vesicle (EV)-associated proteins. Our results highlight the complexity of the apoplastic immune response, where proteins function in distinct stages of PTI activation, defense reinforcement, and signaling. Notably, exosome-associated proteins such as tetraspanin-8 (TET8) were enriched fourfold, and exosome particles increased by approximately threefold. The integration of proteomic and transcriptomic data revealed that transcriptional trends do not always align with protein abundance. This research advances our understanding of the dynamic changes occurring in the apoplast during PTI and underscores the multi-faceted strategies plants employ to defend against pathogen attack. These findings provide a valuable foundation for future investigations into the molecular mechanisms of PTI.

## Materials and Methods

### Plant Tissue Preparation

*A. thaliana* Col-0 seeds suspended in sterile 0.1% agarose were sown in SunGrow 3B Professional potting mix and stratified in darkness for 2 day at 4°C before being grown in a growth chamber (Conviron A1000) with 14-h light (70 μmol m ² s ¹) at 22°C. After 4 weeks, plants were transferred to a growth room maintained under a 12-hour light/12-hour dark cycle for acclimatization. At 4.5 weeks, plants were treated for 16 hours to induce pattern-triggered immunity (PTI). Treatments were applied using a 1 mL blunt-end syringe to 4-5 adult leaves per plant. The treatments consisted of 1 μM flg22 peptide to induce PTI and 0.1% dimethyl sulfoxide (DMSO) as a mock control treatment. The solutions were carefully infiltrated into the abaxial side of the selected leaves, and plants were maintained under the same growth room conditions during the 16-hour treatment period.

### Extraction of Apoplastic Washing Fluid and Cytoplasmic Contamination Measurement

Apoplastic washing fluid (AWF) was crude extracted using vacuum infiltration as described by (Lovelace et al., 2022). 130-150 *A. thaliana* leaves were cut and placed into a 500 mL beaker filled with iced-cold distilled water to top. Repeated cycles of vacuum at 95 kPa for 2 min followed by slow release of pressure were applied until leaves were fully infiltrated. Excess water was blotted from plant tissue before rolled into Saran wrap which were placed into 50 mL conical tubes. Tubes were centrifuged at 1,000 xg for 10 min at 4°C and the fractions were pooled and stored at −80°C. Cytoplasmic contamination in our AWF samples was examined by comparisons of cytosolic marker glucose-6-phosphate dehydrogenase-G6PDH activity in sampled AWF extracts to the total leaf extracts using standard kit (Sigma-Aldrich, St. Louis, MO, USA). Leaves were homogenized at 4°C. G6PDH activity was assayed spectrophotometrically every 5 min at 37°C where each reaction contained 50 μL of AWF and the activity was calculated according to manufacturer’s instructions (Sigma-Aldrich, St. Louis, MO, USA).

### Sample Preparation and Mass Spectrometry Analysis - LC-MS/MS

Samples were filtered through a 3 kDa Amicon spin filter to approximately 50 µL. The retentate was collected via reverse centrifugation, washed with 50 µL of 50 mM Ammonium bicarbonate, and combined. Total protein content was measured using the Pierce BCA Protein Assay Kit (ThermoFisher Scientific) and quantified at 550 nm based on a BSA standard curve. For digestion, 100 µg of protein from each sample was prepared with the EasyPep Mini MS Sample Prep Kit (ThermoFisher Scientific). After reduction and alkylation at 95°C, samples were digested with Trypsin/LysC (0.2 µg/µL) at 37°C for 2 hours. Digestion was stopped, contaminants removed using peptide cleanup columns, and the eluate was dried and resuspended in 3% acetonitrile/0.1% formic acid. Peptide concentration was measured at 205 nm on a NanoDrop, calculated using an extinction coefficient (Scopes, 1974). Reverse-phase chromatography was performed with water + 0.1% formic acid (A) and 80% acetonitrile + 0.1% formic acid (B). A total of 1 µg of peptides was enriched using a PepMap Neo C18 trap-column, followed by separation on a Vanquish Neo system (Thermo Scientific) with a C18 nanospray column at 45°C using a 90 minute gradient at a flow rate of 300 nanoliters/min: 1-6%B over 3 minutes followed by 6-35%B over 70 minutes, 35-45%B over 5 minutes ending in 12 minutes of washing at 500 nanoliters/minute, 99%B. Peptides were eluted directly into an Orbitrap Eclipse mass spectrometer with a Nanospray Flex ion source and analyzed in Data Dependent Acquisition mode. MS spectra were collected over m/z 375–2000 in positive mode, with ions of charge state +2 or higher selected for MS/MS. Dynamic exclusion was set to 1 MS/MS per m/z with a 60 s exclusion. MS detection was performed in FT mode at 240,000 resolution, and MS/MS in ion trap mode with HCD collision energy at 30%.

### Data Analysis and Statistics

Data processing was conducted in Proteome Discoverer (PD) 3.0 (Thermo Scientific). A precursor detector node (S/N=1.5) was used to identify additional precursors within the isolation window for chimeric spectra. Sequest HT searched spectra with methionine oxidation as a dynamic modification and cysteine carbamidomethylating as a fixed modification. An intensity-based rescoring (INFERYS node) using deep learning predicted fragment ion intensities (Zolg et al., 2021). Data were matched against the *Arabidopsis thaliana* Uniprot proteome (UP000006548) and cRAP contaminants. Searches used a fragment ion tolerance of 0.60 Da and parent ion tolerance of 10 PPM. Peptide spectrum matches (PSMs) were validated by Percolator with FDR ≤1% and protein IDs confirmed by at least one peptide. Normalized abundance was log2 transformed. Log2 fold changes (FC) identified proteins with higher (Log2 FC ≥ 1) or lower (Log2 FC ≤ −1) abundance in flg22 vs. mock, with significance tested using t-tests.

### Gene Ontology (GO) Analysis

To identify functional categories enriched in the proteomic data, both significantly enriched proteins and significantly depleted proteins were analyzed using the Gene Ontology (GO) analysis <u>The Arabidopsis Information Resource (TAIR) GO Term Enrichment tool version 2</u>. The analysis was conducted using the TAIR10 genome as the reference dataset to determine over- and under-representation of specific biological processes, molecular functions, and cellular components. Enriched GO terms were considered significant based on a Bonferroni-corrected p-value cutoff of <0.05. The results provided insight into functional trends within the identified apoplastic proteome, including processes associated with plant defense and signal transduction.

### Transcriptomics analysis

To investigate the transcriptional profiles of *Arabidopsis thaliana* proteins identified as significantly enriched or significantly lower abundant in response to flg22 treatment, Tag-Seq data for wild-type Col-0 plants were downloaded from Gene Expression Omnibus (GEO, accession number GSE78735) as described by Hillmer et al., 2017. This dataset comprised transcriptome profiles collected at seven time points post flg22 treatment (0, 1, 2, 3, 5, 9, and 18 hours post-infiltration) with three biological replicates per time point. For each gene, read counts from the three biological replicates were averaged at each time point, normalized to the 0-hour time point by calculating fold changes, and log10-transformed. These processed values were visualized as a heatmap using the pheatmap function from the pheatmap package (version 1.0.12) in R.

### Topology analysis

To analyze the topological distribution of receptor-like kinases (RLKs) in the high-abundance protein profile in response to flg22/mock treatment, peptides identified via Proteome Discoverer (PD) 3.0 were mapped to specific domains or positions within each RLK. Protein domain annotations for the RLKs were obtained from UniProt. Identified peptides corresponding to RLKs were selected and extracted, and their positions were aligned to the respective RLK sequences. This mapping allowed for the determination of peptide coverage within specific protein domains, including extracellular, transmembrane, and intracellular regions.

### Plant Extracellular vesicle isolation and quantification

The plant extracellular vesicles (P-EVs) isolation protocol was modified from previous studies (Huang et al., 2021). Briefly, the crude AWF was centrifuged for 30 min at 4 °C at 2,000 × g to remove large cell debris, and then the supernatant was further centrifuged at 10,000 × g for 30 minutes at 4 °C to remove large insoluble particles. The supernatant (the clean AWF) was then collected and further processed by ultracentrifugation at 100,000 xg for 1 hour at 4 °C (Optima™ TLX Ultracentrifuge, Beckman Coulter, Indianapolis, IN) to purify and concentrate P-EVs. Each P-EV pellet was gently resuspended in 50 µL of 0.22nm-filtered PBS, and all samples were aliquoted and stored at −80 °C until further use. Nano-flow cytometry (NFC, NanoFCM, China) was used to analyze P-EVs isolated from both flg22 and mock samples, following the manufacturer’s instructions. For each sample, a 0.2 µL aliquot of P-EVs was diluted in 50 µL of filtered PBS (1:250 dilution) for analysis. Particle concentration and size distribution were measured in triplicates. To evaluate differences in P-EVs origin between the flg22 and mock groups, yield was calculated as the number of EVs per gram of leaf tissue used for AWF collection. Statistical analysis was performed using Student’s t-test to determine P-EV yield differences between groups (*P<0.05, **P<0.01, NS: not significant).

P-EVs from mock treatment samples were stained with CFSE dye and examined under confocal microscope. The fluorescence signals were observed by using Zeiss LSM 880 confocal laser-scanning microscopy. FITC filter (Excitation: 460–500 nm; Emission: 510–560 nm) (Zeiss, Germany) was used to detect the GFP signal (Excitation: 530–560 nm) (Nikon, Tokyo, Japan). Images were generated and merged using The Zen 2.3 imaging software.

### Data Statement

The mass spectrometry proteomics data is deposited to the ProteomeXchange Consortium via the PRIDE partner repository (Perez-Riverol et al., 2025) with the dataset identifier PXD060654 and 10.6019/PXD060654. Details regarding experimental design, sample preparation, and LC-MS/MS parameters are described in the Materials and Methods section of this manuscript. Total identified proteins, significantly high-abundance proteins by flg22 treatment, and significantly low abundance by flg22 treatment are listed in the Supplementary Data.

## Acknowledgements

This work was supported by the National Science Foundation Grant 1844861 to BK. CJN, LY, and FK was supported by NIH R35GM143067. YZ and YY were supported by startup funds from the University of Georgia (to YY). We thank Su-Yun Uhm from Department of Genetics, University of Georgia for assistance of Arabidopsis planting and AWF collection. We acknowledge the use of resources provided by the Colorado State University ARC-BIO (Research Resource ID: SCR_021758). Data was generated using the Orbitrap Eclipse Mass Spectrometer supported by NSF Grant 2117943. We acknowledge the contribution of ARC-BIO staff-Dorathea Lee for sample preparation.

## Author contributions

HCC, YY, and BHK designed the research; HCC, FK, YZ conducted experiments; HCC, GD, CJN, YZ analyzed the data; HCC, FK, YZ, and BHK wrote the paper. All listed authors reviewed and approved draft and final versions of the manuscript.

## Supporting figures

**Figure S1.**
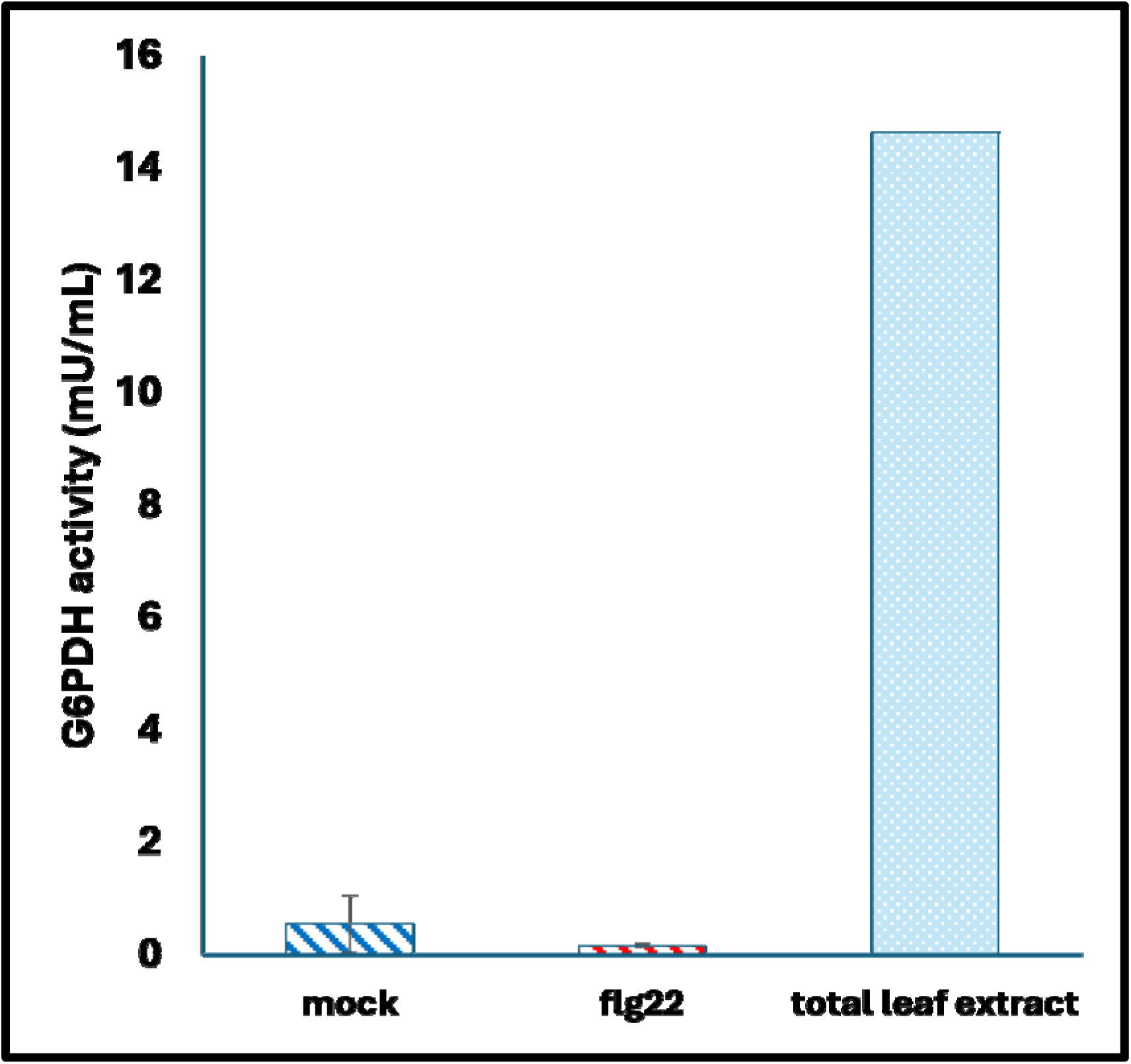
Cytoplasmic contamination detection of apoplastic washing fluid. Glucose-6-phosphate dehydrogenase (G6PDH) enzymatic activity of AWF was tested for cytoplasmic contamination in mock (blue stripe) or flg22 (red stripe) treatment compared to macerated sample (solid). Data are presented as mean ± SD (n=3). Statistical analysis was performed using a two-tailed unpaired Student’s t-test. No significant difference was observed between mock and flg22 treatments (P = 0.256).

**Figure S2.**
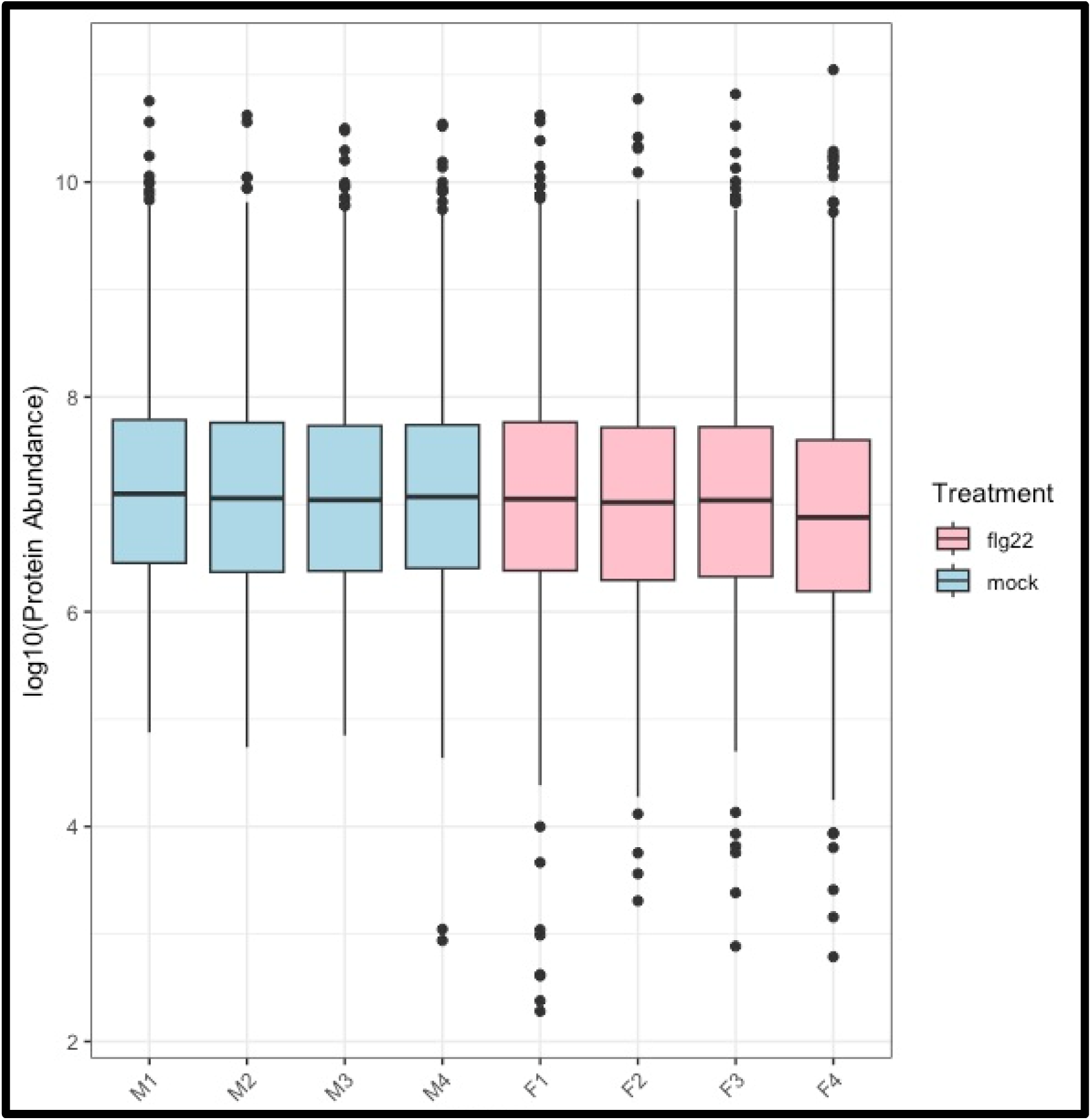
Normalized protein abundances (Log10 transformed data) per sample. Mock treatments are represented in blue boxes (M1-M4) and flg22 treatments are represented in pink boxes (F1-F4). The y-axis represents the log10-transformed protein abundance. Box plots show the distribution of protein abundances, with the box representing the interquartile range (IQR), the horizontal line inside the box indicating the median, and whiskers extending to 1.5 times the IQR. Outliers beyond the whiskers are depicted as individual points.

## References

Acharya BR, Raina S, Maqbool SB, Jagadeeswaran G, Mosher SL, Appel HM, Schultz JC, Klessig DF, Raina R (2007) Overexpression of CRK13, an Arabidopsis cysteine-rich receptor-like kinase, results in enhanced resistance to *Pseudomonas syringa*e. The Plant Journal 50: 488–499

Agrawal GK, Jwa N-S, Lebrun M-H, Job D, Rakwal R (2010) Plant secretome: Unlocking secrets of the secreted proteins. PROTEOMICS 10: 799–827

Alexandersson E, Ali A, Resjö S, Andreasson E (2013) Plant secretome proteomics. Front Plant Sci 4: 9

Anderson JC, Wan Y, Kim Y-M, Pasa-Tolic L, Metz TO, Peck SC (2014) Decreased abundance of type III secretion system-inducing signals in Arabidopsis mkp1 enhances resistance against *Pseudomonas syringae*. Proc Natl Acad Sci U S A 111: 6846–6851

Aung K, Jiang Y, He SY (2018) The role of water in plant–microbe interactions. The Plant Journal 93: 771–780

Bethke G, Grundman RE, Sreekanta S, Truman W, Katagiri F, Glazebrook J (2014) Arabidopsis PECTIN METHYLESTERASEs Contribute to Immunity against *Pseudomonas syringae*. Plant Physiology 164: 1093–1107

Bigeard J, Colcombet J, Hirt H (2015) Signaling Mechanisms in Pattern-Triggered Immunity (PTI). Molecular Plant 8: 521–539

Boavida LC, Qin P, Broz M, Becker JD, McCormick S (2013) Arabidopsis tetraspanins are confined to discrete expression domains and cell types in reproductive tissues and form homo-and heterodimers when expressed in yeast. Plant Physiol 163: 696–712

Boller T, He SY (2009) Innate immunity in plants: an arms race between pattern recognition receptors in plants and effectors in microbial pathogens. Science 324: 742–744

Cai Q, He B, Weiberg A, Buck AH, Jin H (2019) Small RNAs and extracellular vesicles: New mechanisms of cross-species communication and innovative tools for disease control. PLOS Pathogens 15: e1008090

Cai Q, Qiao L, Wang M, He B, Lin F-M, Palmquist J, Huang S-D, Jin H (2018) Plants send small RNAs in extracellular vesicles to fungal pathogen to silence virulence genes. Science 360: 1126–1129

Camejo D, Guzmán-Cedeño Á, Moreno A (2016) Reactive oxygen species, essential molecules, during plant-pathogen interactions. Plant Physiol Biochem 103: 10–23

Chan C, Panzeri D, Okuma E, Tõldsepp K, Wang Y-Y, Louh G-Y, Chin T-C, Yeh Y-H, Yeh H-L, Yekondi S, Huang Y-H, Huang T-Y, Chiou T-J, Murata Y, Kollisit H, Zimmerli L (2020) STRESS INDUCED FACTOR 2 Regulates Arabidopsis Stomatal Immunity through Phosphorylation of the Anion Channel SLAC1. Plant Cell 32: 2216–2236

DeFalco TA, Zipfel C (2021) Molecular mechanisms of early plant pattern-triggered immune signaling. Molecular Cell 81: 3449–3467

Delaunois B, Colby T, Belloy N, Conreux A, Harzen A, Baillieul F, Clément C, Schmidt J, Jeandet P, Cordelier S (2013) Large-scale proteomic analysis of the grapevine leaf apoplastic fluid reveals mainly stress-related proteins and cell wall modifying enzymes. BMC Plant Biology 13: 24

Delaunois B, Jeandet P, Clément C, Baillieul F, Dorey S, Cordelier S (2014) Uncovering plant-pathogen crosstalk through apoplastic proteomic studies. Front Plant Sci. doi: 10.3389/fpls.2014.00249

Djamei A, Pitzschke A, Nakagami H, Rajh I, Hirt H (2007) Trojan Horse Strategy in Agrobacterium Transformation: Abusing MAPK Defense Signaling. Science 318: 453–456

Farvardin A, González-Hernández AI, Llorens E, García-Agustín P, Scalschi L, Vicedo B (2020) The Apoplast: A Key Player in Plant Survival. Antioxidants (Basel) 9: 604

Figueiredo L, Santos RB, Figueiredo A (2021) Defense and Offense Strategies: The Role of Aspartic Proteases in Plant–Pathogen Interactions. Biology (Basel) 10: 75

Fiorin GL, Sanchéz-Vallet A, Thomazella DP de T, Prado PFV do, Nascimento LC do, Figueira AV de O, Thomma BPHJ, Pereira GAG, Teixeira PJPL (2018) Suppression of Plant Immunity by Fungal Chitinase-like Effectors. Current Biology 28: 3023–3030.e5

Flinn BS (2008) Plant extracellular matrix metalloproteinases. Functional Plant Biol 35: 1183–1193

Fonseca JP, Dong X (2014) Functional Characterization of a Nudix Hydrolase AtNUDX8 upon Pathogen Attack Indicates a Positive Role in Plant Immune Responses. PLOS ONE 9: e114119

Freeman BC, Beattie GA (2009) Bacterial Growth Restriction During Host Resistance to *Pseudomonas syringae* Is Associated with Leaf Water Loss and Localized Cessation of Vascular Activity in *Arabidopsis thaliana*. MPMI 22: 857–867

Gao C, Zhao Y, Wang W, Zhang B, Huang X, Wang Y, Tang D (2024) BRASSINOSTEROID-SIGNALING KINASE 1 modulates OPEN STOMATA 1 phosphorylation and contributes to stomatal closure and plant immunity. The Plant Journal 120: 45–59

Gentzel I, Giese L, Ekanayake G, Mikhail K, Zhao W, Cocuron J-C, Alonso AP, Mackey D (2022) Dynamic nutrient acquisition from a hydrated apoplast supports biotrophic proliferation of a bacterial pathogen of maize. Cell Host & Microbe 30: 502–517.e4

Hillmer RA, Tsuda K, Rallapalli G, Asai S, Truman W, Papke MD, Sakakibara H, Jones JDG, Myers CL, Katagiri F (2017) The highly buffered Arabidopsis immune signaling network conceals the functions of its components. PLoS Genet 13: e1006639

Huang Y, Wang S, Cai Q, Jin H (2021) Effective methods for isolation and purification of extracellular vesicles from plants. J Integr Plant Biol 63: 2020–2030

Huot B, Yao J, Montgomery BL, He SY (2014) Growth–Defense Tradeoffs in Plants: A Balancing Act to Optimize Fitness. Molecular Plant 7: 1267–1287

Jung Y-H, Jeong S-H, Kim SH, Singh R, Lee J, Cho Y-S, Agrawal GK, Rakwal R, Jwa N-S (2008) Systematic Secretome Analyses of Rice Leaf and Seed Callus Suspension-Cultured Cells: Workflow Development and Establishment of High-Density Two-Dimensional Gel Reference Maps. J Proteome Res 7: 5187–5210

Kaffarnik FAR, Jones AME, Rathjen JP, Peck SC (2009) Effector Proteins of the Bacterial Pathogen *Pseudomonas syringae* Alter the Extracellular Proteome of the Host Plant, *Arabidopsis thaliana*. Molecular & Cellular Proteomics 8: 145–156

Kim ST, Kang YH, Wang Y, Wu J, Park ZY, Rakwal R, Agrawal GK, Lee SY, Kang KY (2009) Secretome analysis of differentially induced proteins in rice suspension-cultured cells triggered by rice blast fungus and elicitor. Proteomics 9: 1302–1313

Liebminger E, Veit C, Pabst M, Batoux M, Zipfel C, Altmann F, Mach L, Strasser R (2011) β-N-Acetylhexosaminidases HEXO1 and HEXO3 Are Responsible for the Formation of Paucimannosidic N-Glycans in *Arabidopsis thaliana*\*. Journal of Biological Chemistry 286: 10793–10802

Liu G, Kang G, Wang S, Huang Y, Cai Q (2021) Extracellular Vesicles: Emerging Players in Plant Defense Against Pathogens. Front Plant Sci. 12: 757925

Liu N, Hou L, Chen X, Bao J, Chen F, Cai W, Zhu H, Wang L, Chen X (2024) Arabidopsis TETRASPANIN8 mediates exosome secretion and glycosyl inositol phosphoceramide sorting and trafficking. The Plant Cell 36: 626–641

Liu Z, Hou S, Rodrigues O, Wang P, Luo D, Munemasa S, Lei J, Liu J, Ortiz-Morea FA, Wang X, et al (2022) Phytocytokine signalling reopens stomata in plant immunity and water loss. Nature 605: 332–339

van Loon LC, Rep M, Pieterse CMJ (2006) Significance of inducible defense-related proteins in infected plants. Annu Rev Phytopathol 44: 135–162

Lovelace AH, Chen H-C, Lee S, Soufi Z, Bota P, Preston GM, Kvitko BH (2022) RpoS contributes in a host-dependent manner to *Salmonella* colonization of the leaf apoplast during plant disease. Front Microbiol 13: 999183

Martínez-González AP, Ardila HD, Martínez-Peralta ST, Melgarejo-Muñoz LM, Castillejo-Sánchez MA, Jorrín-Novo JV (2018) What proteomic analysis of the apoplast tells us about plant–pathogen interactions. Plant Pathology 67: 1647–1668

McLellan H, Gilroy EM, Yun B-W, Birch PRJ, Loake GJ (2009) Functional redundancy in the Arabidopsis Cathepsin B gene family contributes to basal defence, the hypersensitive response and senescence. New Phytologist 183: 408–418

Mott GA, Middleton MA, Desveaux D, Guttman DS (2014) Peptides and small molecules of the plant-pathogen apoplastic arena. Front Plant Sci. doi: 10.3389/fpls.2014.00677

Munzert KS, Engelsdorf T (2025) Plant cell wall structure and dynamics in plant–pathogen interactions and pathogen defence. Journal of Experimental Botany 76: 228–242

Nemati M, Singh B, Mir RA, Nemati M, Babaei A, Ahmadi M, Rasmi Y, Golezani AG, Rezaie J (2022) Plant-derived extracellular vesicles: a novel nanomedicine approach with advantages and challenges. Cell Communication and Signaling 20: 69

Nishimura M (2016) Cell wall reorganization during infection in fungal plant pathogens. Physiological and Molecular Plant Pathology 95: 14–19

O’Leary BM, Neale HC, Geilfus C, Jackson RW, Arnold DL, Preston GM (2016) Early changes in apoplast composition associated with defence and disease in interactions between *Phaseolus vulgaris* and the halo blight pathogen *Pseudomonas syringae* pv. *phaseolicola*. Plant Cell Environ 39: 2172–2184

Perez-Riverol Y, Bandla C, Kundu DJ, Kamatchinathan S, Bai J, Hewapathirana S, John NS, Prakash A, Walzer M, Wang S, Vizcaino JA. (2025) The PRIDE database at 20 years: 2025 update. Nucleic Acids Res 53: D543–D553

Perrot T, Pauly M, Ramírez V (2022) Emerging Roles of β-Glucanases in Plant Development and Adaptative Responses. Plants (Basel) 11: 1119

Ramírez V, López A, Mauch-Mani B, Gil MJ, Vera P (2013) An Extracellular Subtilase Switch for Immune Priming in Arabidopsis. PLOS Pathogens 9: e1003445

Roussin-Léveillée C, Mackey D, Ekanayake G, Gohmann R, Moffett P (2024) Extracellular niche establishment by plant pathogens. Nat Rev Microbiol 22: 360–372

Rubiato HM, Liu M, O’Connell RJ, Nielsen ME (2022) Plant SYP12 syntaxins mediate an evolutionarily conserved general immunity to filamentous pathogens. eLife 11: e73487

Rutter BD, Innes RW (2017) Extracellular Vesicles Isolated from the Leaf Apoplast Carry Stress-Response Proteins. Plant Physiol 173: 728–741

Scopes RK (1974) Measurement of protein by spectrophotometry at 205 nm. Analytical Biochemistry 59: 277–282

Serag A, Salem MA, Gong S, Wu J-L, Farag MA (2023) Decoding Metabolic Reprogramming in Plants under Pathogen Attacks, a Comprehensive Review of Emerging Metabolomics Technologies to Maximize Their Applications. Metabolites 13: 424

Torres MA, Jones JDG, Dangl JL (2006) Reactive oxygen species signaling in response to pathogens. Plant Physiol 141: 373–378

Van Der Hoorn RAL (2008) Plant Proteases: From Phenotypes to Molecular Mechanisms. Annu Rev Plant Biol 59: 191–223

Vlot AC, Sales JH, Lenk M, Bauer K, Brambilla A, Sommer A, Chen Y, Wenig M, Nayem S (2021) Systemic propagation of immunity in plants. New Phytologist 229: 1234–1250

Waghmare S, Lileikyte E, Karnik R, Goodman JK, Blatt MR, Jones AME (2018) SNAREs SYP121 and SYP122 Mediate the Secretion of Distinct Cargo Subsets1. Plant Physiol 178: 1679–1688

Wang J, Sun W, Kong X, Zhao C, Li J, Chen Y, Gao Z, Zuo K (2020) The peptidyl-prolyl isomerases FKBP15-1 and FKBP15-2 negatively affect lateral root development by repressing the vacuolar invertase VIN2 in Arabidopsis. Planta 252: 52

Wang Z, Zeng J, Deng J, Hou X, Zhang J, Yan W, Cai Q (2023) Pathogen-Derived Extracellular Vesicles: Emerging Mediators of Plant-Microbe Interactions. MPMI 36: 218–227

Xin X-F, Nomura K, Aung K, Velásquez AC, Yao J, Boutrot F, Chang JH, Zipfel C, He SY (2016) Bacteria establish an aqueous living space in plants crucial for virulence. Nature 539: 524–529

Yadeta KA, Elmore JM, Creer AY, Feng B, Franco JY, Rufian JS, He P, Phinney B, Coaker G (2017) A Cysteine-Rich Protein Kinase Associates with a Membrane Immune Complex and the Cysteine Residues Are Required for Cell Death1. Plant Physiol 173: 771–787

Yeh Y-H, Panzeri D, Kadota Y, Huang Y-C, Huang P-Y, Tao C-N, Roux M, Chien H-C, Chin T-C, Chu P-W, Zipfel C, Zimmerli L (2016) The Arabidopsis Malectin-Like/LRR-RLK IOS1 Is Critical for BAK1-Dependent and BAK1-Independent Pattern-Triggered Immunity. The Plant Cell 28: 1701–1721

Zhang C, He M, Jiang Z, Liu L, Pu J, Zhang W, Wang S, Xu F (2022) The Xyloglucan Endotransglucosylase/Hydrolase Gene XTH22/TCH4 Regulates Plant Growth by Disrupting the Cell Wall Homeostasis in Arabidopsis under Boron Deficiency. Int J Mol Sci 23: 1250

Zhao P, Zhang F, Liu D, Imani J, Langen G, Kogel K-H (2017) Matrix metalloproteinases operate redundantly in Arabidopsis immunity against necrotrophic and biotrophic fungal pathogens. PLoS One 12: e0183577

Zolg DP, Gessulat S, Paschke C, Graber M, Rathke-Kuhnert M, Seefried F, Fitzemeier K, Berg F, Lopez-Ferrer D, Horn D, Henrich C, Huhmer A, Delanghe B, Frejno M (2021) INFERYS rescoring: Boosting peptide identifications and scoring confidence of database search results. Rapid Communications in Mass Spectrometry n/a: e9128

